## supplementary figures and tables for "Pre-stimulus beta power mediates explicit and implicit perceptual biases in distinct cortical areas"

### Forster et al.

#### Supplementary figures:

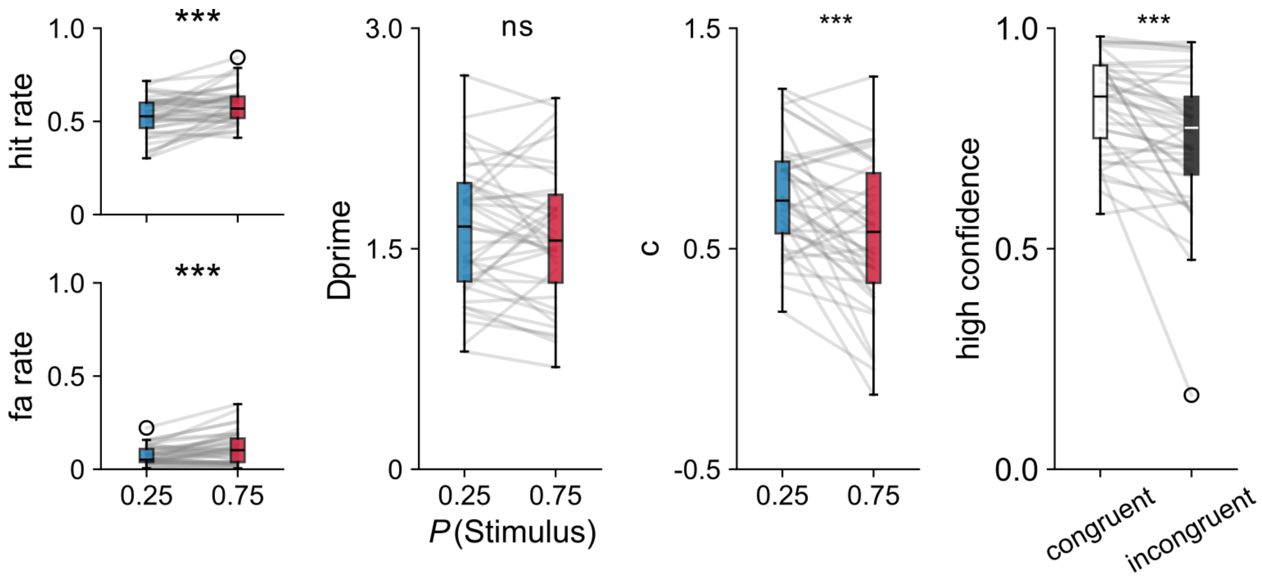


**Fig. 1.1: Behavioural results after excluding participants with a false alarm rate > 40 % (n=3) in high expectation condition.**

All statistical results remain after the exclusion. Box plots depict median and interquartile ranges, while whiskers show minimum and maximum values. **Significance levels:** *** $p < .001$, **$p < .01$, *$p < .05$. **Abbreviations:** ns = not significant.

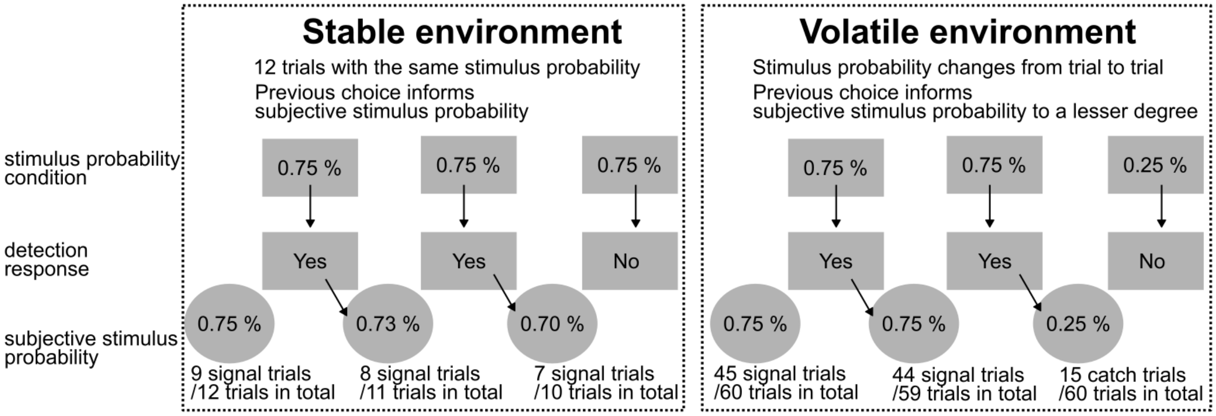


**Fig. 1.2: The previous choice has a greater influence on the subjective stimulus probability in the stable compared to the volatile environment.** The block design creates small environments of a stable stimulus probability. In the example (left box) the stimulus probability is high (0.75 % of all trials contain a stimulus in the block). An ideal human observer tries to track the previous choices and updates the current (subjective) stimulus probability accordingly. The previous choice has a lesser influence on the subjective stimulus probability in the trial-by-trial design (right box) as the participant must track the previous choices over the whole experimental block of 120 trials.

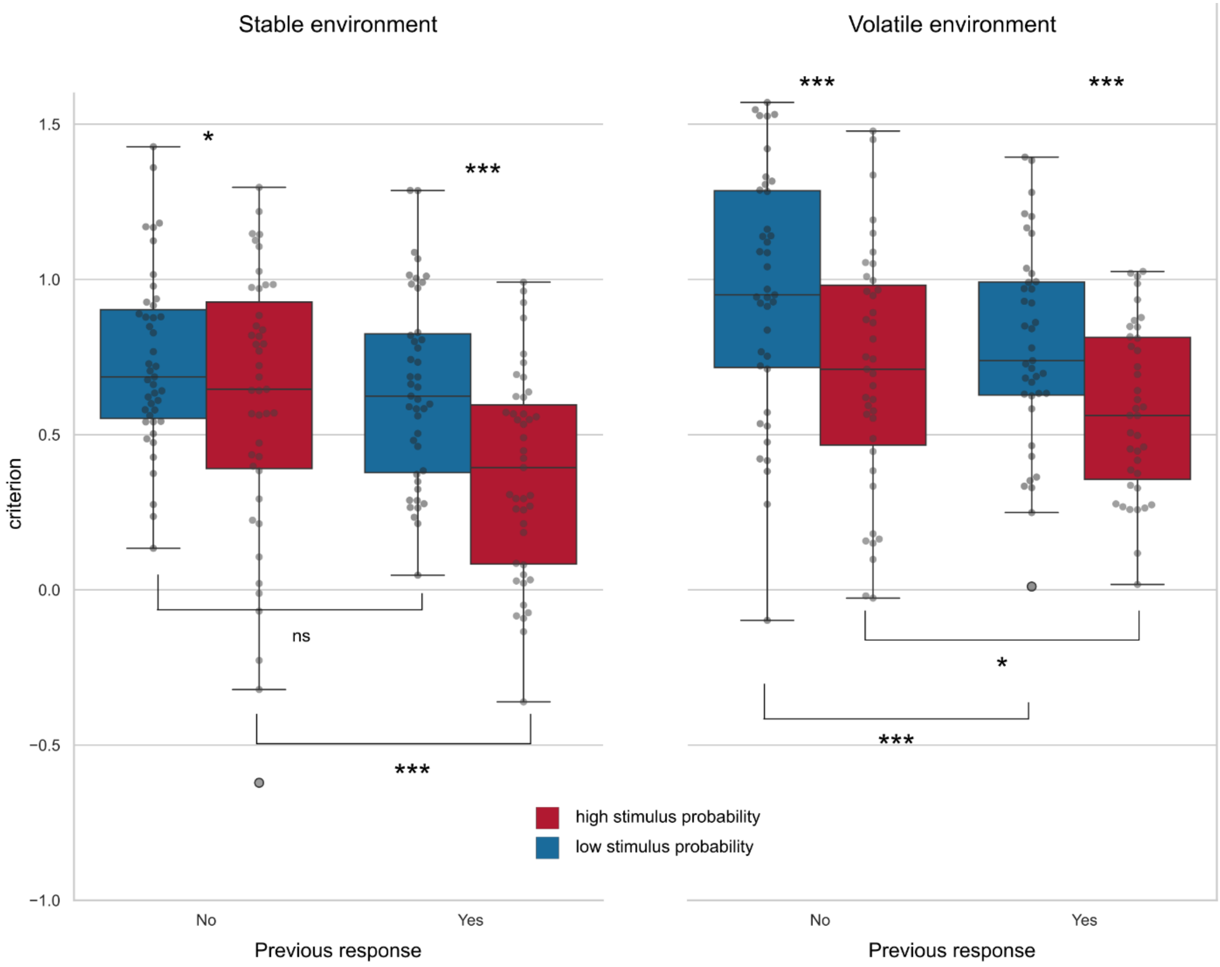


**Fig. 1.3: Model-free analysis of the interaction between criterion and previous response:** The criterion is always significantly higher (more conservative) in the low probability condition in both environments. The criterion is also always more conservative after previous no responses in both environments, except for the low probability condition in the stable environment. This is in line with the model-based results. Box plots depict median and interquartile ranges, while whiskers show minimum and maximum values. **Significance levels: *** ** $p < .001$, ** ** ** $p < .01$, ** * ** $p < .05$. ns = not significant.

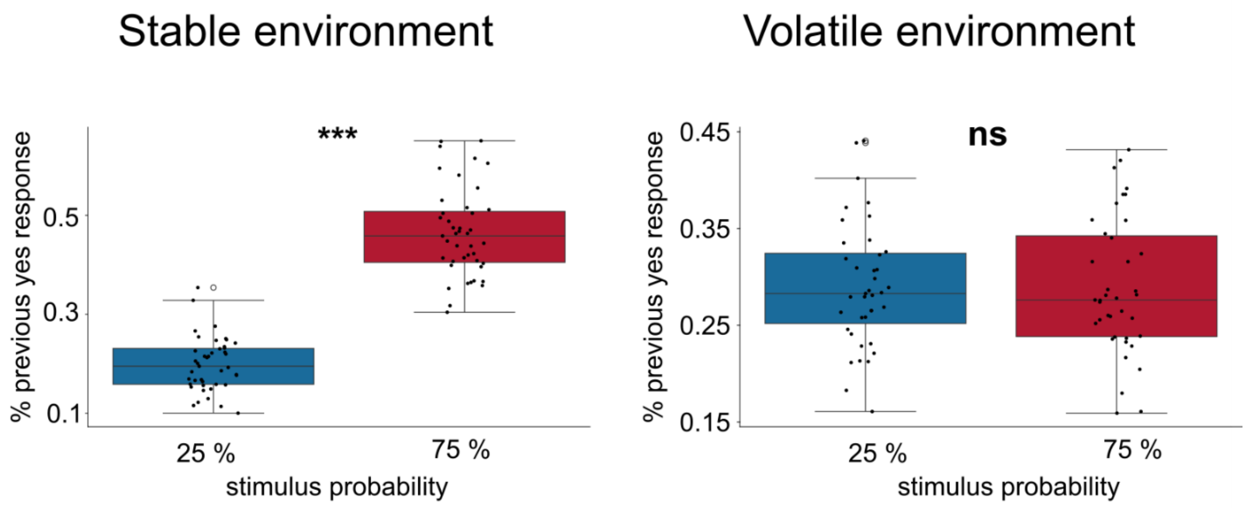


**Fig. 1.4**: **Previous response distributions differ significantly in the stable but not in the volatile environment.** There are significantly fewer previous yes responses in the low probability condition in the stable environment, but there is no significant difference in the distribution of previous responses for the volatile environment due to the randomisation of probability cues in the second study. Box plots depict median and interquartile ranges, while whiskers show minimum and maximum values. **Significance levels: ***** $p < .001$, ****** $p < .01$, ***** $p < .05$. ns = not significant.

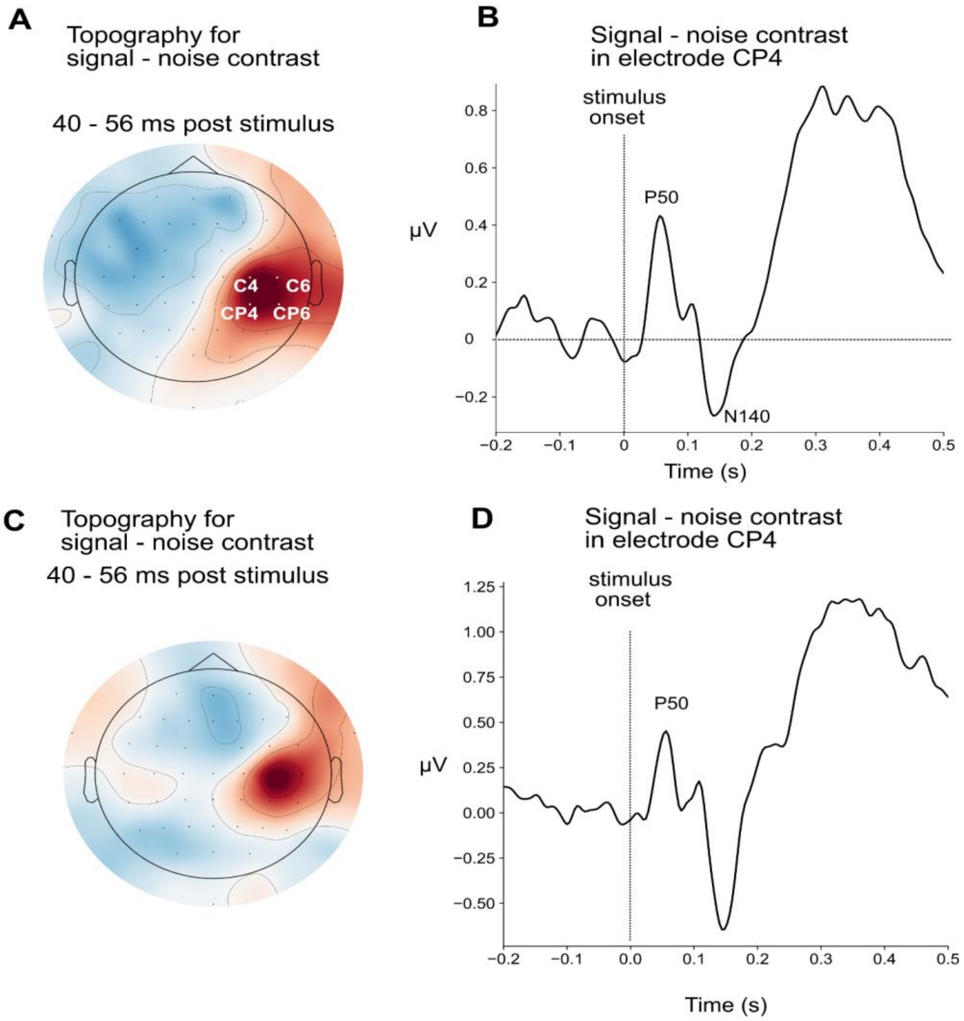


**Fig. 2.1: Somatosensory region of interest definition based on the contrast between signal and noise trials. A:** Topography for the contrast signal-noise around the earliest somatosensory evoked potential (P50), averaged between 40 and 56 ms in the stable environment. **B:** Somatosensory evoked potential averaged over participants for the contrast signal-noise trials in electrode CP4 in the volatile environment. **C:** Topography for the contrast signal - noise around the earliest somatosensory evoked potential (P50), averaged between 40 and 56 ms in the stable environment. **D:** Somatosensory evoked potential averaged over participants for the contrast signal-noise trials in electrode CP4 in the volatile environment. The signal was baseline corrected with a mean from -100ms to stimulus onset.

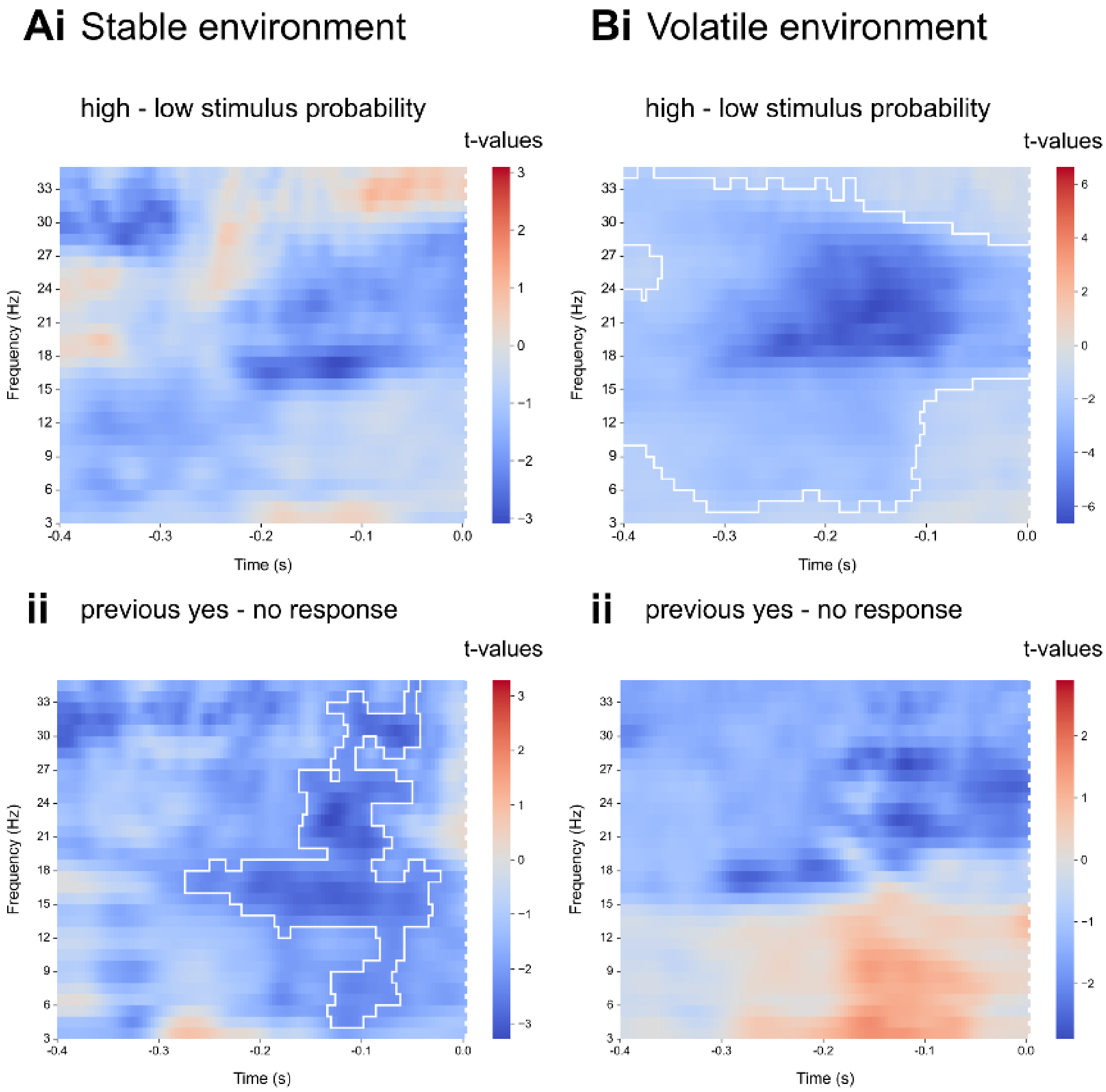


**Fig. 2.2: Cluster-based permutation test for a shorter pre-stimulus window.** Time-frequency contrasts for high–low stimulus probability (first row) and previous yes - no response (second row). The time window was restricted to 400ms. Significant clusters (cluster p-value < .05 are highlighted with a white border).

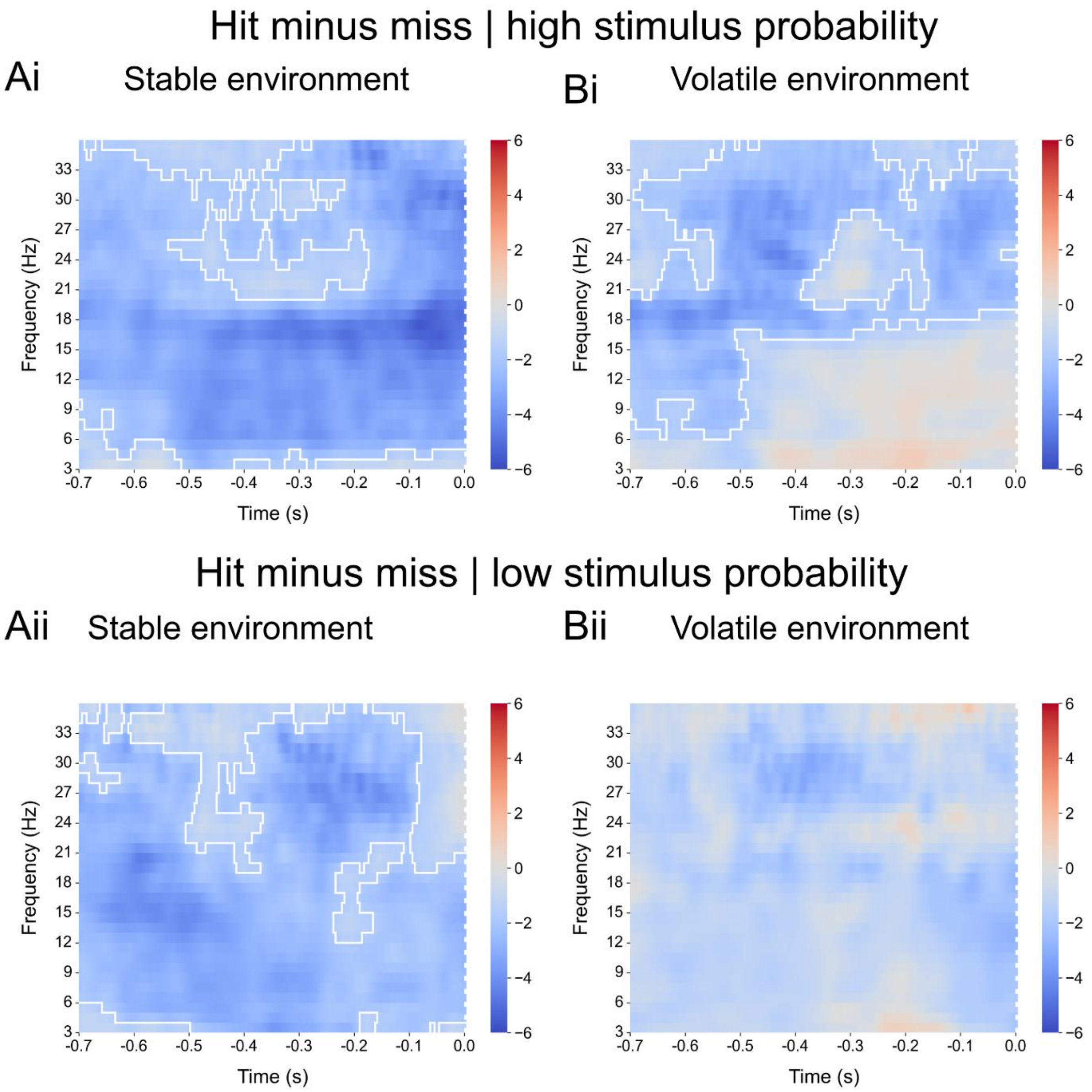


**Fig. 2.3: Time-Frequency contrast hits minus misses in electrode CP4: Ai:** Threshold-free cluster-based permutation test in low frequencies in the pre-stimulus window for hits (detected signals) minus misses(undetected signals) contrast in the high probability condition shows a significant cluster ($p < .001$) with lower power in alpha and beta frequencies in the stable probability environment. **Bi:** In the volatile environment, the cluster t. ($p < .001$) spans alpha and beta frequencies after cue offset, but only beta frequencies shortly before stimulus onset. **Aii:** The cluster test for the hits minus misses contrast in the low probability condition shows a significant cluster ($p < .001$) in the stable probability environment, with lower alpha and beta power before hits. **Bii:** There is no significant cluster (minimum $p = .063$) in the volatile probability environment for hits minus misses in the low stimulus probability condition. Significant clusters are highlighted with a white border.

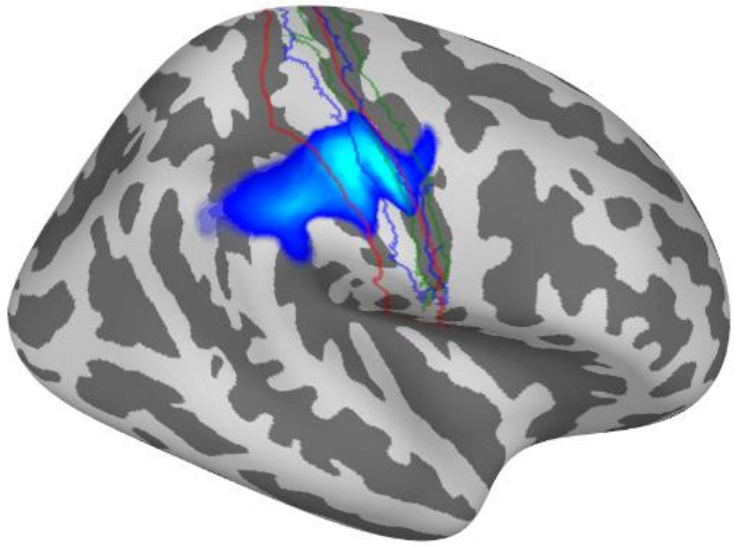

**Fig. 2.4: Control analysis for source reconstruction using DICS beamforming:** Beta power source reconstruction for signal–noise contrast in the post-stimulus window. Lighter blue values indicate more negative t-values. The area highlighted in red marks the postcentral gyrus.

#### Supplementary tables:

**Table 1:** Generalised linear mixed effects model predicting the detection response on each trial in the stable environment.

**Stable environment**
**Outcome variable: detection response**

| | **Base** estimate, 95 % confidence interval, p-value | **add probability** estimate, 95 % confidence interval, p-value | **add previous response** estimate, 95 % confidence interval, p-value | **Interaction** estimate, 95 % confidence interval, p-value |
|---|---|---|---|---|
| (Intercept) | -1.437 [-1.570, -1.305], *** | -1.518 [-1.647, -1.389], *** | -1.573 [-1.711, -1.435], *** | -1.546 [-1.684, -1.408], *** |
| Stimulus [1] | 1.637 [1.506, 1.767], *** | 1.584 [1.438, 1.730], *** | 1.607 [1.456, 1.758], *** | 1.599 [1.449, 1.749], *** |
| Probability [0.75] | | 0.259 [0.157, 0.360], *** | 0.193 [0.089, 0.297], *** | 0.141 [0.034, 0.248], ** |
| Stimulus × Probability | | -0.079 [-0.162, 0.005], + | -0.076 [-0.160, 0.009] | -0.065 [-0.150, 0.019], |
| Previous response [1] | | | 0.231 [0.149, 0.312], *** | 0.117 [0.018, -0.216], * |
| Prob. x prev. resp. | | | | 0.163 [0.080, 0.245], *** |
| SD (Intercept ID) | 0.428, | 0.409, | 0.440, | 0.439, |
| SD (prob. ID) | | 0.250, | 0.257, | 0.254, |
| SD (stimulus ID) | 0.416, | 0.449, | 0.466, | 0.463, |
| SD (prev. resp. ID) | | | 0.239, | 0.235, |
| Num. Obs. | 28776 | 28776 | 28776 | 28776 |
| R2 Marg. | 0.375 | 0.381 | 0.387 | 0.387 |
| R2 Cond. | 0.440 | 0.451 | 0.462 | 0.461 |
| AIC | 27547.9 | 27318.0 | 27098.1 | 27085.2 |
| BIC | 27589.2 | 27400.6 | 27222.1 | 27217.5 |
| ICC | 0.1 | 0.1 | 0.1 | 0.1 |
| RMSE | 0.40 | 0.39 | 0.39 | 0.39 |

*** $p < .001$, ** $p < .01$, * $p < .05$

**Table 2** Generalised linear mixed effects model predicting the detection response on each trial in the volatile environment.

**Volatile environment**
**Outcome variable: detection response**

| | **Base** estimate, 95 % confidence interval, p-value | **add probability** estimate, 95 % confidence interval, p-value | **add previous response** estimate, 95 % confidence interval, p-value | **Interaction** estimate, 95 % confidence interval, p-value |
|---|---|---|---|---|
| (Intercept) | -1.743 [-1.921, -1.565], *** | -1.796 [-1.977, -1.615], *** | -1.863 [-2.066, -1.659], *** | -1.866 [-2.070, -1.661], *** |
| Stimulus [1] | 1.853 [1.668, 2.038], *** | 1.742 [1.543, 1.942], *** | 1.764 [1.559, 1.970], *** | 1.765 [1.560, 1.970], *** |
| Probability [0.75] | | 0.174 [0.066, 0.283], ** | 0.180 [0.071, 0.289], ** | 0.184 [0.071, 0.298], ** |
| Stimulus × Prob. | | 0.043 [-0.067, 0.153], | 0.040 [-0.070, 0. 151], | 0.040 [-0.070, 0.150], |
| Previous response [1] | | | 0.177 [0.092, 0.261], *** | 0.185 [0.082, 0.288], *** |
| Probability x Prev. resp. | | | | -0.014 [-0.108, 0.081], |
| SD (Intercept ID) | 0.540, | 0.541, | 0.613, | 0.613, |
| SD (stimulus ID) | 0.558, | 0.583, | 0.602, | 0.603, |
| SD (probability ID) | | 0.168, | 0.169, | 0.169, |
| SD (prev. resp. ID) | | | 0.222, | 0.222, |
| Num. Obs. | 21421 | 21421 | 21421 | 21421 |
| R2 Marg. | 0.420 | 0.423 | 0.427 | 0.427 |
| R2 Cond. | 0.511 | 0.516 | 0.527 | 0.527 |
| AIC | 18834.3 | 18747.9 | 18666.9 | 18668.8 |
| BIC | 18874.2 | 18827.6 | 18786.5 | 18796.4 |
| ICC | 0.2 | 0.2 | 0.2 | 0.2 |
| RMSE | 0.38 | 0.38 | 0.38 | 0.38 |

*** p < .001, ** p < .01, * p < .05

**Table 3** Generalised linear mixed effects model predicting the detection response in each trial, including beta power in the stable environment.

**Stable environment**
**Outcome variable: detection response**

| | **Base**<br>estimate, 95 % confidence interval, p-value | **add previous response**<br>estimate, 95 % confidence interval, p-value | **Interaction**<br>estimate, 95 % confidence interval, p-value |
|---|---|---|---|
| (Intercept) | -1.526 [-1.654, -1.398], *** | -1.579 [-1.717, -1.442], *** | -1.555 [-1.694, -1.417], *** |
| Stimulus [1] | 1.578 [1.432, 1.725], *** | 1.601 [1.450, 1.753], *** | 1.594 [1.444, 1.744], *** |
| Pre-stimulus beta | -0.074 [-0.129, -0.018], ** | -0.068 [-0.123, -0.012], * | -0.099 [-0.159, -0.039], ** |
| Probability [0.75] | 0.258 [0.156, 0.360], *** | 0.193 [0.089, 0.297], *** | 0.141 [0.034, 0.248], ** |
| Stimulus × beta | -0.010 [-0.076, 0.057], | -0.007 [-0.073, 0.059], | -0.005 [-0.071, 0.062], |
| Stimulus × Prob. | -0.071 [-0.156, 0.014], | -0.068 [-0.153, 0.017], | -0.058 [-0.143, 0.028], |
| Previous response | | 0.225 [0.144, 0.306], *** | 0.119 [0.018, 0.220], * |
| Beta × prev. resp. | | | 0.073 [0.017, 0.129], * |
| Prob. × prev. resp. | | | 0.164 [0.081, 0.248], *** |
| SD (Intercept ID) | 0.407, | 0.438, | 0.438, |
| SD (isyes1 ID) | 0.447, | 0.465, | 0.462, |
| SD (cue0.75 ID) | 0.250, | 0.255, | 0.253, |
| SD (prevresp1 ID) | | 0.239, | 0.240, |
| Num. Obs. | 28290 | 28290 | 28290 |
| R2 Marg. | 0.383 | 0.389 | 0.389 |
| R2 Cond. | 0.454 | 0.464 | 0.464 |
| AIC | 26853.0 | 26644.3 | 26626.7 |
| BIC | 26952.0 | 26784.5 | 26783.5 |
| ICC | 0.1 | 0.1 | 0.1 |
| RMSE | 0.39 | 0.39 | 0.39 |

**Table 4** Generalised linear mixed effects model predicting the detection response in each trial, including beta power in the volatile environment.

**Volatile environment**
**Outcome variable: detection response**

| | **Base** estimate, 95 % confidence interval, p-value | **add previous response** estimate, 95 % confidence interval, p-value | **Interaction** estimate, 95 % confidence interval, p-value |
|---|---|---|---|
| (Intercept) | -1.822 [-2.086, -1.638], *** | -1.882 [-2.086, -1.678], *** | -1.886 [-2.092, -1.680], *** |
| Stimulus [1] | 1.763 [1.565, 1.961], *** | 1.783 [1.580, 1.986], *** | 1.784 [1.581, 1.988], *** |
| Pre-stimulus Beta | -0.192 [-0.268, -0.117], *** | -0.175 [-0.249, -0.101], *** | -0.195 [-0.276, -0.114], *** |
| Probability [0.75] | 0.152 [0.042, 0.261], ** | 0.158 [0.048, 0.268], ** | 0.160 [0.045, 0.274], ** |
| Stimulus x beta | 0.073 [-0.013, 0.159], + | 0.063 [-0.021, 0.148], | 0.072 [-0.014, 0.158], |
| Stimulus x prob. | 0.057 [-0.055, 0.169], | 0.054 [-0.058, 0.166], | 0.054 [-0.058, 0.167], |
| Previous response | | 0.164 [0.082, 0.246], *** | 0.175 [0.072, 0.278], *** |
| Beta x prev. resp. | | | 0.036 [-0.025, 0.098], |
| Prob. × prev. resp. | | | -0.007 [-0.103, 0.089], |
| SD (Intercept ID) | 0.548, | 0.612, | 0.616, |
| SD (stimulus ID) | 0.573, | 0.591, | 0.592, |
| SD (prob. ID) | 0.163, | 0.166, | 0.166, |
| SD (prev. resp. ID) | | 0.211, | 0.214, |
| Num. Obs. | 20903 | 20903 | 20903 |
| R2 Marg. | 0.431 | 0.434 | 0.435 |
| R2 Cond. | 0.528 | 0.536 | 0.537 |
| AIC | 18253.9 | 18187.3 | 18190.0 |

| | **Base**<br><br>estimate, 95 % confidence interval, p-value | **add previous response**<br>estimate, 95 % confidence interval, p-value | **Interaction**<br>estimate, 95 % confidence interval, p-value |
|---|---|---|---|
| BIC | 18349.3 | 18322.5 | 18341.0 |
| ICC | 0.2 | 0.2 | 0.2 |
| RMSE | 0.38 | 0.38 | 0.38 |

*** $p < .001$, ** $p < .01$, * $p < .05$

**Table 5** Generalised linear mixed effects model predicting the confidence response in each trial, including either the probability cue or pre-stimulus beta power in the stable environment.

**Stable environment**

**Outcome variable: confidence rating**

| | **Probability**<br><br>estimate, 95 % confidence interval, p-value | **Beta power**<br><br>estimate, 95 % confidence interval, p-value |
|---|---|---|
| (Intercept) | 0.831 [0.600, 1.062], *** | 0.519 [0.271, 0.766], *** |
| Response [1] | -1.229 [-1.434, -1.023], *** | -0.897 [-1.096, -0.698], *** |
| Probability [0.75] | -0.443 [-0.585, -0.300], *** | |
| Previous confidence | 0.184 [0.089, 0.280], *** | 0.209 [0.109, 0.309], *** |
| Accuracy [correct] | 0.597 [0.521, 0.673], *** | 0.744 [0.665, 0.823], *** |
| Response × Probability | 0.673 [0.456, 0.889], *** | |
| Pre-stimulus beta power | | 0.084 [0.039, 0.129], *** |
| Response × Beta | | -0.106 [-0.174, -0.038], ** |
| SD (Intercept ID) | 0.729, | 0.802, |
| SD (response ID) | 0.647, | 0.647, |
| SD (accuracy ID) | 0.187, | 0.211, |
| SD (previous confidence ID) | 0.260, | 0.279, |
| SD (probability ID) | 0.427, | |
| SD (response x cue ID) | 0.654, | |
| Num. Obs. | 28290 | 28290 |
| R2 Marg. | 0.166 | 0.148 |
| R2 Cond. | 0.430 | 0.399 |
| AIC | 24019.5 | 24535.0 |
| BIC | 24242.3 | 24667.0 |
| ICC | 0.3 | 0.3 |
| RMSE | 0.37 | 0.37 |

*** $p < .001$, ** $p < .01$, * $p < .05$

**Table 6** Generalised linear mixed effects model predicting the confidence response in each trial, including either the probability cue or pre-stimulus beta power in the volatile environment.

**Volatile environment**

**Outcome variable: confidence rating**

| | **Probability** estimate, 95 % confidence interval, p-value | **Beta power** estimate, 95 % confidence interval, p-value |
|---|---|---|
| (Intercept) | 0.754 [0.496, 1.012], *** | 0.657 [0.380, 0.934], *** |
| Response [1] | -1.301 [-1.513, -1.089], *** | -1.133 [-1.371, -0.895], *** |
| Probability [0.75] | -0.147 [-0.259, -0.034], * | |
| Previous confidence | 0.226 [0.122, 0.330], *** | 0.226 [0.121, 0.331], *** |
| Accuracy [correct] | 0.627 [0.518, 0.737], *** | 0.695 [0.588, 0.802], *** |
| Response × Probability | 0.342 [0.177, 0.506], *** | |
| Pre-stimulus beta power | | 0.045 [-0.006, 0.096], |
| Response × Beta | | -0.095 [-0.175, -0.016], * |
| SD (Intercept ID) | 0.755, | 0.838, |
| SD (response ID) | 0.601, | 0.720, |
| SD (accuracy ID) | 0.243, | 0.251, |
| SD (previous confidence ID) | 0.224, | 0.228, |
| SD (probability ID) | 0.235, | |
| SD (response x cue ID) | 0.352, | |
| Num. Obs. | 20903 | 20903 |
| R2 Marg. | 0.155 | 0.153 |
| R2 Cond. | 0.482 | 0.485 |
| AIC | 17926.9 | 18005.0 |
| BIC | 18141.5 | 18132.1 |
| ICC | 0.4 | 0.4 |
| RMSE | 0.37 | 0.37 |

*** p < .001, ** p < .01, * p < .05

**Table 7** Generalised linear mixed effects models for mediation analysis predicting the detection response by including pre-stimulus beta power for the probability contrast or pre-stimulus beta power for the previous response contrast in the stable environment.

**Stable environment**
**Outcome variable: detection response**

| | **Probability** estimate, 95 % confidence interval, p-value | **Previous response** estimate, 95 % confidence interval, p-value |
|---|---|---|
| (Intercept) | -1.541 [-1.677, -1.405], *** | -1.548 [-1.680, -1.415], *** |
| Beta_power_prob | -0.071 [-0.103, -0.040], *** | |
| Previous response [1] | 0.108 [0.009, 0.207], * | 0.082 [-0.015, 0.180], |
| Stimulus probability [0.75] | 0.101 [0.010, 0.191], * | 0.140 [0.027, 0.253], * |
| Stimulus [1] | 1.569 [1.423, 1.714], *** | 1.590 [1.441, 1.738], *** |
| Prev. resp × Probability | 0.168 [0.085, 0.252], *** | 0.192 [0.090, 0.293], *** |
| Beta_power_prev | | -0.061 [-0.092, -0.030], *** |
| Probability x Stimulus | | -0.055 [-0.140, 0.031], |
| SD (Intercept ID) | 0.435, | 0.418, |
| SD (stimulus ID) | 0.464, | 0.455, |
| SD (previous response ID) | 0.234, | 0.205, |
| SD (probability ID) | 0.256, | 0.280, |
| Num. Obs. | 28280 | 28280 |
| R2 Marg. | 0.388 | 0.388 |
| R2 Cond. | 0.462 | 0.463 |
| AIC | 26621.9 | 26630.3 |
| BIC | 26753.9 | 26811.8 |
| ICC | 0.1 | 0.1 |
| RMSE | 0.39 | 0.39 |

*** $p < .001$, ** $p < .01$, * $p < .05$

**Table 8** Generalised linear mixed effects models for mediation analysis predicting the detection response by including pre-stimulus beta power for the probability contrast or pre-stimulus beta power for the previous response contrast in the volatile environment.

**Volatile environment**
**Outcome variable: detection response**

| | **Probability** estimate, 95 % confidence interval, p-value | **Previous response** estimate, 95 % confidence interval, p-value |
|---|---|---|
| (Intercept) | -1.883 [-2.085, -1.682], *** | -1.878 [-2.079, -1.677], *** |
| Beta power prob. | -0.128 [-0.166, -0.090], *** | |
| Previous response [1] | 0.164 [0.082, 0.246], *** | 0.166 [0.084, 0.249], *** |
| Stimulus probability [0.75] | 0.198 [0.126, 0.270], *** | 0.203 [0.131, 0.275], *** |
| Stimulus [1] | 1.793 [1.597, 1.989], *** | 1.790 [1.594, 1.987], *** |
| Beta power prev. | | -0.078 [-0.118, -0.039], *** |
| SD (Intercept ID) | 0.610, | 0.609, |
| SD (stimulus ID) | 0.587, | 0.589, |
| SD (previous response ID) | 0.211, | 0.213, |
| SD (probability ID) | 0.165, | 0.167, |
| Num.Obs. | 20892 | 20892 |
| R2 Marg. | 0.433 | 0.430 |
| R2 Cond. | 0.534 | 0.530 |
| AIC | 18182.8 | 18211.8 |
| BIC | 18302.1 | 18331.0 |
| ICC | 0.2 | 0.2 |
| RMSE | 0.38 | 0.38 |

*** $p < .001$, ** $p < .01$, * $p < .05$
